# MetaFusion: A high-confidence metacaller for filtering and prioritizing RNA-seq gene fusion candidates

**DOI:** 10.1101/2020.09.17.302307

**Authors:** Michael Apostolides, Yue Jiang, Mia Husić, Robert Siddaway, Cynthia Hawkins, Andrei L. Turinsky, Michael Brudno, Arun K. Ramani

## Abstract

**Motivation:** Current fusion detection tools use diverse calling approaches and provide varying results, making selection of the appropriate tool challenging. Ensemble fusion calling techniques appear promising; however, current options have limited accessibility and function.

**Results:** MetaFusion is a flexible meta-calling tool that amalgamates outputs from any number of fusion callers. Individual caller results are standardized by conversion into the new file type Common Fusion Format (CFF). Calls are annotated, merged using graph clustering, filtered, and ranked to provide a final output of high confidence candidates. MetaFusion consistently achieves higher precision and recall than individual callers on real and simulated datasets, and reaches up to 100% precision, indicating that ensemble calling is imperative for high confidence results. MetaFusion uses FusionAnnotator to annotate calls with information from cancer fusion databases, and is provided with a benchmarking toolkit to calibrate new callers.

**Availability:** MetaFusion is freely available at https://github.com/ccmbioinfo/MetaFusion

**Supplementary information:** Supplementary data are available at Bioinformatics online.

## 1. Introduction

Gene fusions, or hybridizations between two independent genes, are recognized as an important class of genomic alterations, particularly in cancer. They arise most frequently from chromosomal rearrangements, though recent evidence indicates that they are also caused by mechanisms such as cis-splicing of adjacent genes (cis-SAGe) (Brien et al. 2019; Gao et al. 2018; Grosso et al. 2015; Hu et al. 2018; Li et al. 2008). Cancer-related fusions can lead to increased oncogene expression, decreased tumour suppressor expression, and formation of oncogenic fusion proteins. Such changes are well-documented in the tumourigenesis of multiple cancers, such as lymphoma, chronic myeloid leukemia (CML), and lung cancer (Mertens et al. 2015; Mitelman et al. 2007; Xiao et al. 2018; Yoshihara et al. 2015), and it is estimated that they occur in nearly 20% of all cancers (Gao et al. 2018; Mitelman et al. 2007). Fusions may serve as disease biomarkers, such as the breast cancer-specific *SCNN1A-TNFRSF1A* and *CTSD-IFITM10* (Varley et al. 2014), or as treatment targets, such as the tyrosine kinase activity of the causative *BCR-ABL* fusion in CML, which is inhibited by imatinib (Druker 2008; Mitelman et al. 2007). Accurate identification of biologically relevant gene fusions in cancer is thus critical, as it can contribute to patient diagnosis and care in the rise of precision medicine.

Although a number of tools are currently available for fusion calling in human cancer samples, they vary significantly from one another. Callers differ in the genomic regions they consider, the numbers of alignment steps they have, read coverage requirements, filters, output formats, and so on. Some may only prioritize specific types of fusions, such as those caused by chromosomal rearrangements, considering all others to be transcriptional noise or part of normal cell biology. Such differences can give inconsistent results between methods for any single dataset and lead to biologically relevant fusions being excluded from final outputs. These problems are further compounded by individual fusion caller limitations, including low precision (high false positive rate) and sub-par recall. Poor precision can be caused by outdated filters, reliance on outdated databases of false positive fusions, or having overly lenient criteria for keeping reads. Likewise, benchmarking with only simulated data may result in more false positive calls than expected when a caller is run on real cancer data. Regarding poor recall, some callers have stringent criteria for keeping reads, and low read depth on a true fusion may cause it to be missed. Often, a tool which performs well in one of these areas will fall short in another (Liu et al. 2016). Finally, fusion callers often provide outputs that are large and ambiguous (Haas et al. 2019), making it a challenge to prioritize biologically relevant fusions for further experimental validation. This leaves users with the arduous task of determining which tools are best suited for their needs.

Similar challenges have been overcome in various fields within genetics and gene expression through the use of ensemble approaches (Aghaeepour et al. 2013; Huang et al. 2019; Lichtenberg et al. 2017; Yang and Deng 2020), and a recent study of 23 fusion callers has indeed shown that using multiple tools leads to improved fusion calling results (Haas et al. 2019). Yet current means of ensemble fusion identification, or meta-calling, have been largely preliminary (Liu et al. 2016) and, to our knowledge, no robust approach has been developed. Software tools such as Pegasus (Abate et al. 2014) may standardize the interface of various callers, but do not merge their outputs effectively, making downstream analysis difficult. Chimera collates results from 10 callers and can visualize junction coverage and predict the oncogenic potential of a given fusion (Beccuti et al. 2014). Yet this is not a stand-alone meta-caller, functioning instead as a library only compatible with the output files of its 10 pre-defined callers, making it limited and inflexible in its utility. Fusion search engine-based approaches such as FusionHub also exist, but harness information in existing databases as opposed to combining the results of various callers (Panigrahi et al. 2018).

These obstacles highlight the need for a single, flexible method that utilizes multiple approaches for fusion identification and evaluation to provide a concise and high-confidence list of candidate fusions. We have thus developed MetaFusion, an ensemble fusion calling method that incorporates predictions from any number of callers, leveraging the results of all the tools included. It takes fusion calls in a Common Fusion Format (CFF), a novel file type that we have developed to standardize fusion caller outputs. MetaFusion uses a series of filters to remove false positives for optimal precision, ranks calls based on the number of contributing tools, and can utilize existing fusion databases to further annotate calls. Fusion caller combinations can be customized according to the user’s needs and tool availability, and can be calibrated using a Benchmarking Toolkit. Using a series of simulated and real cancer datasets, we show that MetaFusion consistently achieves high precision and recall, and provides high-confidence candidate fusions for experimental validation in research or clinical contexts. As it can incorporate any number of fusion callers, it is highly adaptable and can be easily updated as new tools become available.

## 2. Methods

### 2.1 MetaFusion workflow

The MetaFusion workflow was developed to consolidate the outputs from various pre-existing fusion callers and allow for further filtering, refining, and benchmarking of those outputs (Figure 1). Metafusion has been developed as a standalone tool, and the workflow to run the fusion callers has been implemented in GenPipes (Bourgey et al. 2019), an open-source, Python-based framework for -omics pipeline development and deployment. The current GenPipes implementation contains seven fusion callers, with MetaFusion downstream from them for consolidation and further analysis of their outputs. MetaFusion dependencies are also available as Docker and Singularity images. MetaFusion dependencies can likewise be installed directly onto one’s machine, although using a container is the preferred method.

**Figure 1.**
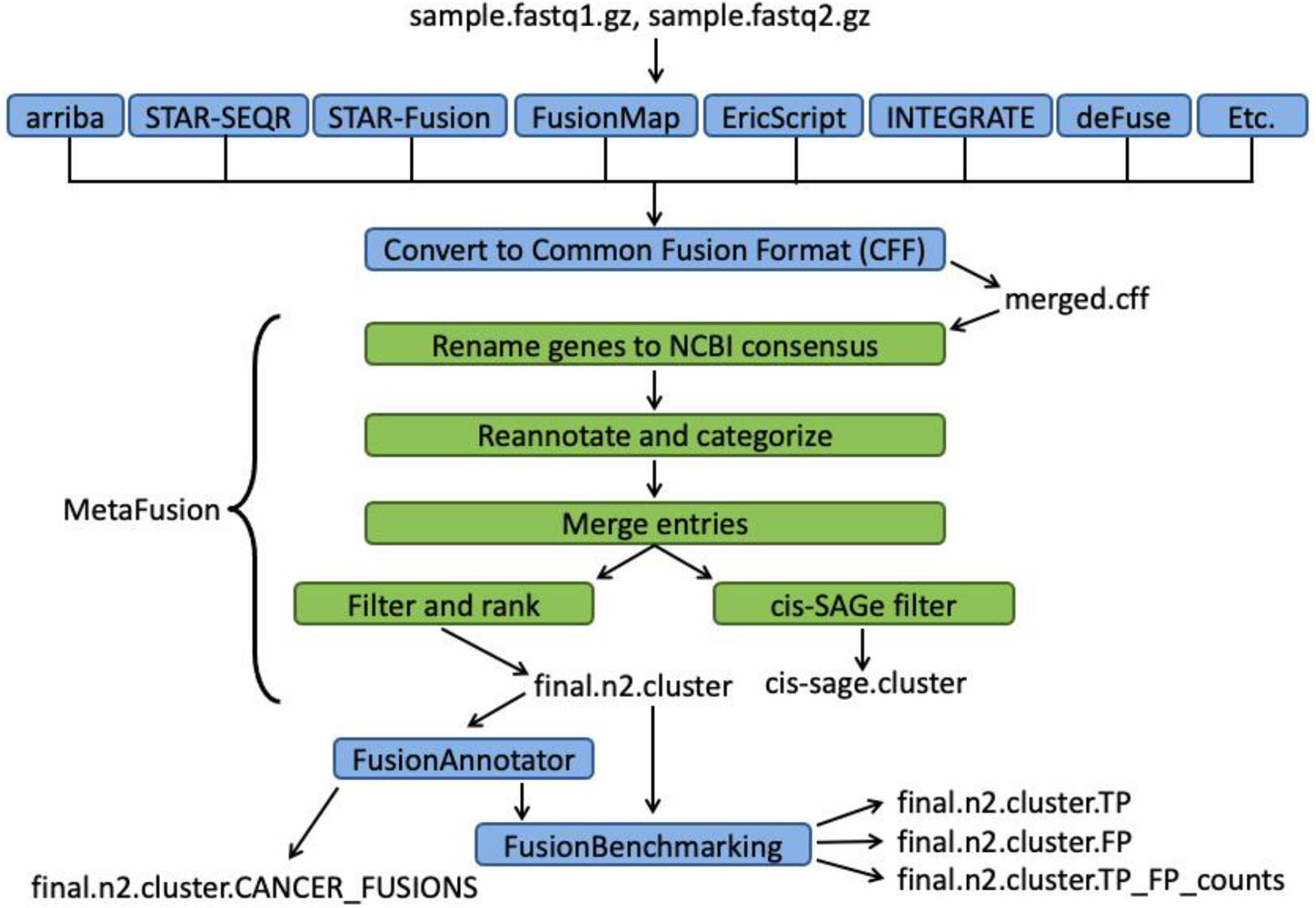
The MetaFusion workflow. MetaFusion consolidates outputs from various fusion callers. The seven callers used in this study are shown, while “Etc.” indicates that any tools the user wishes to include may be incorporated in addition to or instead of these seven. The callers are run independently on a given dataset and their outputs are converted into CFF files, which are used as input into the MetaFusion pipeline (green). This consists of renaming, reannotation, categorization, merging and filtering steps. Afterwards, tools such as FusionAnnotator and the Benchmarking Toolkit may be used to further refine the results.

We measured the running time and memory requirements of MetaFusion on a local machine. 7GB of RAM was required, and running time was linear with the number of input calls and can range from one to two minutes in instances with a few hundred calls, to 30 minutes in instances with over 20,000 calls. Such a relatively short running time, even with large inputs, makes MetaFusion straightforward and practical to incorporate into fusion analysis workflows.

#### 2.1.1 Fusion calling and caller output conversion to CFF

For this study, we chose seven fusion callers to use as input into MetaFusion: deFuse (McPherson et al. 2011), Arriba (Uhrig 2020), STAR-SEQR (STAR-SEQR 2020), STAR-Fusion (Haas et al. 2017), INTEGRATE (Zhang et al. 2016), EricScript (Benelli et al. 2012), and FusionMap (Ge et al. 2011). These callers are widely used in the literature and represent a broad array of fusion calling approaches, allowing us to capture an array of fusion calling strategies upstream of MetaFusion. Furthermore, STAR-SEQR, Arriba, and STAR-Fusion were among the top-performing callers in a recent, large-scale benchmarking study of fusion calling tools (Haas et al. 2019). Each caller is independently run on the input data to generate its own fusion calls.

All seven callers provide outputs in differing formats. To standardize this, we have developed the CFF file format (example shown in Supplementary Table 1). Prior to the start of the MetaFusion workflow, a wrapper script converts fusion caller outputs into CFF. Separate sections of this script exist for each caller, where caller output file fields are mapped to CFF fields. Each line in a CFF file represents one fusion call by a given caller (Supplementary Table 1). Subsequent MetaFusion steps add further information to the CFF file, such as a unique fusion identifier (FID; e.g. F00000001) and fusion category (see Methods below).

While we chose the callers mentioned above any number and combination of fusion callers can be used with the MetaFusion pipeline. Users can modify the wrapper script to include new callers, or convert to CFF using a method of their choosing. This means that users have the flexibility to easily incorporate tools outside of those selected here as input into the MetaFusion pipeline. As a demonstration, we provide an use-case examples with an eighth caller, ChimeraScan (Iyer et al. 2011), and a smaller subset of four callers (ChimeraScan, INTEGRATE, EricScript and deFuse) to detect cis-SAGe ReadThrough fusions in prostate cancer (Supplementary Figure 1). These four callers were chosen as they detect the highest number of true positive cis-SAGe fusions.

#### 2.1.2 Renaming of genes to current NCBI symbols

Renaming genes to current NCBI symbols optimizes subsequent merging and benchmarking steps in the MetaFusion workflow that rely on matching gene names. This is done using the NCBI *Homo sapiens* gene alias file (*Homo_sapiens*.*gene_info*.*gz*, accessed May 7 2020), which is freely available on the NCBI FTP website (ftp://ftp.ncbi.nlm.nih.gov/gene). If the gene name is neither a known NCBI symbol nor an alias of one, the original gene name is retained.

#### 2.1.3 Reannotation and categorization

Once all gene names are updated to current NCBI symbols, each fusion entry in the CFF is reannotated according to the following:

1. Each CFF entry is assigned a unique fusion identifier (FID; e.g. F00000001)
2. Breakpoints for each fusion are reannotated based on their intersection with genomic features (e.g. exon, intron, 5’UTR, 3’UTR, etc) in the gene annotation file. If multiple intersections occur for a given breakpoint, the genomic feature that matches the gene name is chosen.
3. Each fusion entry in the CFF is assigned to one of seven categories based on the coding status and adjacency of the fusion partners (Figure 2). For any given fusion, the upstream gene is referred to as the “head gene” and the downstream gene is the “tail gene.” A fusion’s annotated category can be used to prioritize and filter fusion candidates based on user needs.

**Figure 2.**
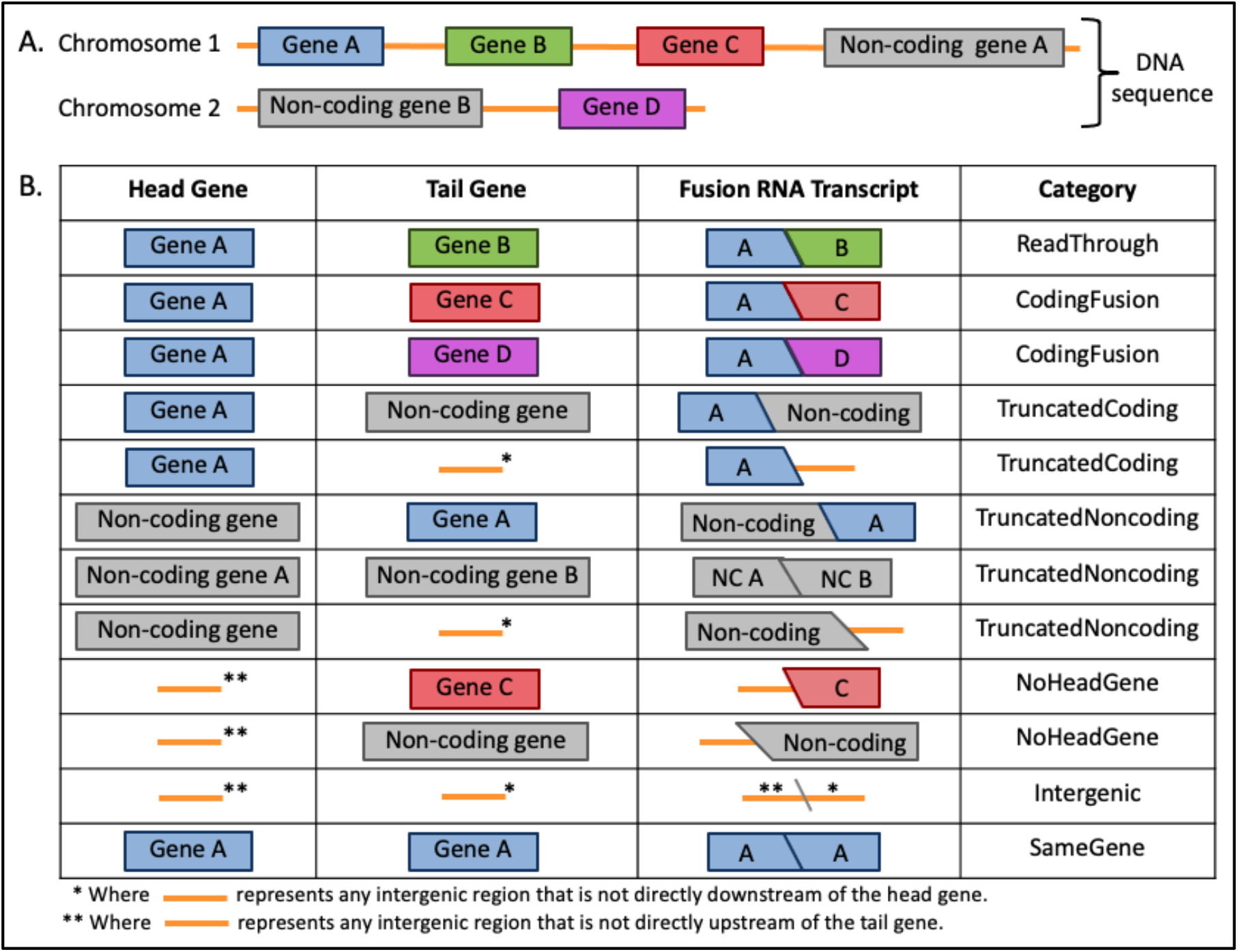
The seven categories used by MetaFusion to distinguish fusions. **(A)** Two chromosomes containing coding (coloured) and non-coding (grey) genes, with intergenic sequences represented by an orange line between genes. **(B)** From the DNA sequences shown in **A**, head and tail genes, the resultant fusion RNA transcripts and the fusion categories that the transcripts would be assigned to. NC: non-coding.

#### 2.1.4 Merging of fusion calls

After reannotation and categorization, fusion calls are merged together using breakpoints and gene names (Figure 3). To consolidate fusion calls from multiple callers, we applied a graph-clustering algorithm, in which nodes represent individual fusion calls from each caller, and edges represent intersections based on breakpoints, gene names, or both. Breakpoints and gene names are considered together to allow for the most complete merging of calls and provide a concise final output. Altogether, this process is done in four steps. See Supplementary Materials for details.

**Figure 3.**
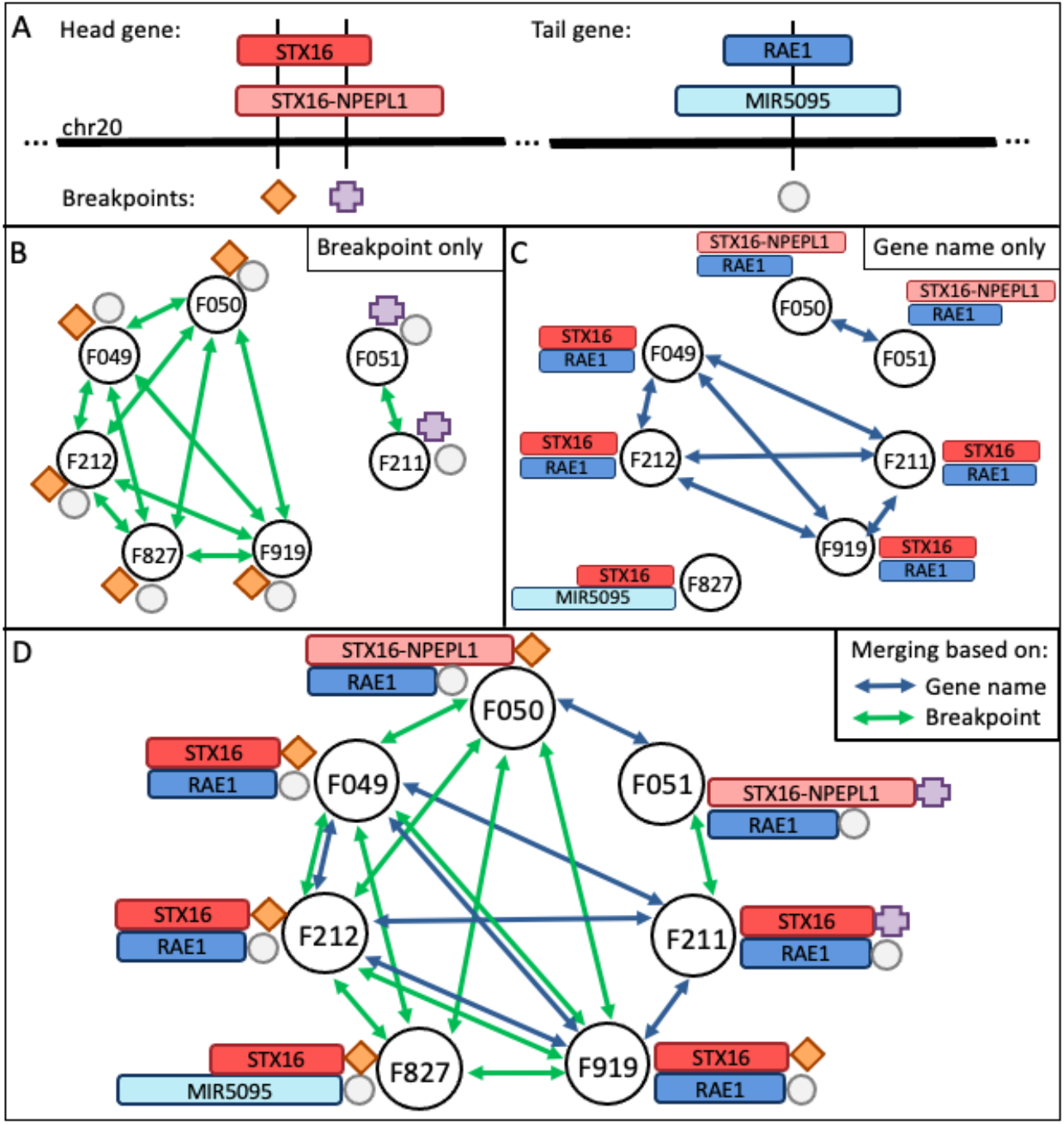
Diagram of the graph clustering approach used to merge calls in MetaFusion. The BT474 breast cancer true positive *STX16-RAE1* fusion is used to show how MetaFusion merges calls based on both breakpoints and gene names to arrive at its final output. **(A)** Schematic of chromosome 20, showing approximate placements of the fusion’s head and tail genes, gene name variants, and the breakpoints identified in each gene. Breakpoints are indicated by the diamond, cross, and circle shapes. In panels **B-D**, corresponding breakpoints are indicated by the same shapes, and edges represent intersections based on breakpoint, gene name, or both. Edge colours indicate if calls are merged based on breakpoint (green) or name (blue). If only breakpoints **(B)** or only genes names **(C)** are considered for merging, calls may be merged incorrectly or orphaned, resulting in excess entries in the final output. **(D)** MetaFusion relies on both breakpoint and gene name intersections to determine if multiple calls should be merged into one. FIDs correspond to those in Supplementary Table 1A, and corresponding CFF can be found in the test data .cff in test_data/cff/BRCA.cff.

#### 2.1.5 Custom filters

After merging of fusion calls is complete, the resulting list is refined using a series of filters. This includes the *blocklist* filter, *callerfilterN, ReadThrough* filter and *adjacent noncoding* filter. Details of the filters is provided in the Supplementary Methods.

#### 2.1.6 Benchmarking Toolkit

The Benchmarking Toolkit (Haas et al. 2019) allows for the benchmarking of MetaFusion outputs. When a caller combination other than the one used here is chosen, MetaFusion should be benchmarked with the provided test data to ensure adequate performance (please see *Software Availability* and Github wiki for FASTQ files). This series of perl scripts has been modified slightly to include the unique FIDs provided by MetaFusion’s reannotation step that allows the MetaFusion output to be partitioned into separate true positive and false positive files. We likewise modified the Benchmarking Toolkit’s *genes*.*coords*.*gz* file to include an additional 41,496 entries corresponding to loci with updated NCBI symbols in NCBI’s most recent *Homo_sapiens*.*gene_info* file (accessed May 7th, 2020). This ensures that all gene naming is consistent with the most up-to-date NCBI symbols.

More information about the Benchmarking Toolkit can be found at https://github.com/fusiontranscripts/FusionBenchmarking/wiki.

#### 2.1.7 FusionAnnotator

As part of the Benchmarking Toolkit, MetaFusion integrates the FusionAnnotator tool (Haas et al. 2019), which leverages several cancer fusion databases to annotate calls with previously seen fusions (see FusionAnnotator documentation for further details). Although run as part of the Benchmarking Toolkit (Step 7), FusionAnnotator functions as a separate step unrelated to benchmarking. It annotates fusion calls with metadata from a number of cancer and normal fusion databases, based on gene name matches. An enrichment in cancer-related fusions may indicate that a workflow is prioritizing fusions of interest. FusionAnnotator output is used to generate “CANCER_FUSION” and “NORMAL_FUSION” subsets of MetaFusion output.

### 2.2 Datasets

To evaluate MetaFusion, we analyzed a series of simulated and real cancer datasets (described below, with additional information available in Supplementary Table 2) using the MetaFusion pipeline. CFF files for all data in this study are available at https://github.com/ccmbioinfo/MetaFusion/tree/master/test_data/cff.

#### DREAM dataset

This simulated dataset comprises the sim45 and sim52 datasets from the SMC DREAM RNA challenge. The sim45 dataset contains 60 million reads, 101bp long with a fragment size of 150-160. The sim52 dataset contains 135 million reads, 101bp long with a fragment size of 150-160.

#### Negative control BEERS dataset

This simulated dataset was created using the Benchmarker for Evaluating the Effectiveness of RNA-seq software (BEERS) simulator (Grant et al. 2011) by the authors of JAFFA (Davidson et al. 2015), and contains no true fusions. This dataset is used to identify fusion callers with high false positive rates, and to determine the characteristics of false positive fusions.

#### sim50 and sim101 datasets

Both the sim50 and sim101 simulated datasets were generated using the Fusion Transcript Simulation Toolkit (Haas et al. 2019). The sim50 dataset has 50 base pair reads. The sim101 dataset has 101 base pair reads.

#### Breast cancer (BRCA)

This dataset consists of the BT474, KPL4, MCF7, and SKBR3 breast cancer cell lines (Edgren et al. 2011). It was previously used as a benchmarking dataset to evaluate the performances of 23 fusion callers (Haas et al. 2019). These samples were downloaded from the Broad Institute’s Trinity index.

#### Melanoma and chronic myeloid leukemia

The melanoma-CML dataset consists of melanoma patient-derived samples (SRR018259 SRR018260 SRR018261 SRR018265 SRR018266 SRR018267) and two CML cell lines (SRR018268, SRR018269) (Supplementary Table 3) (Berger et al. 2010; Jia et al. 2013). It was previously used for benchmarking the SOAPfuse fusion caller (Jia et al. 2013).

#### NTRK

This dataset consists of 15 *NTRK* (neurotrophic tyrosine receptor kinase) fusions with GM24385 human genomic RNA as background (Seracare). The 15 fusions in the truth set contain 12 unique head-tail gene pairs, which we use for our benchmarking. Unlike the samples in the BRCA and melanoma-CML datasets, which were sequenced using whole RNA-seq, this dataset was sequenced using the TruSight pan-cancer panel (Illumina), which searches for fusions in 1385 fusion-associated genes.

#### Prostate cancer

This dataset primarily contains ReadThrough fusions (Kumar et al. 2016; Qin et al. 2015). This data is divided into 100nt read length (SRR1657556, SRR1657557) and 50nt read length (SRR1657558 SRR1657559, SRR1657560 and SRR1657561). Samples SRR1657557, SRR1657559 and SRR1657561 are siCTCF-treated, whereas SRR1657556, SRR1657558 and SRR1657560 are negative controls. This dataset was chosen for a use case showing how various callers can be used with MetaFusion.

#### 2.2.1 Curation and re-naming of truth sets

Evaluation of fusion calling methods can be challenging due to a lack of clearly defined truth sets. These are subsets of fusions that have been either intentionally created in simulated data or experimentally confirmed in real cancer data. Several of the above datasets were selected due to their previous use in benchmarking of fusion calling methods (Haas et al. 2019; Jia et al. 2013). As the Benchmarking Toolkit relies on gene name matches to identify true and false positives, it is important that the names of genes involved in these truth set fusions follow NCBI consensus naming. Although some benchmarking approaches might use breakpoints, this can be a challenge as it may involve unannotated regions of the genome where it is difficult to distinguish noise from biologically relevant events.

Upon renaming the truth set for sim50/101, NCBI symbols were updated for at least one fusion partner in 257/2500 fusions. For this reason, we have used the NCBI *Homo_sapiens*.*gene_info* file to update names to the most recent NCBI symbols in the truth sets. Truth sets were run through the renaming script in a separate step, independent of the MetaFusion workflow.

## 3. Results

### 3.1 Precision and recall of MetaFusion and individual callers

To evaluate MetaFusion’s performance, we compared its precision and recall to that of the seven individual callers we used. We examined the performance of MetaFusion with all seven callers, as well using only STAR-SEQR, Arriba, and STAR-Fusion (MetaFusion.top_3), which were shown to be top performers among fusion calling tools in a recent benchmarking study (Haas et al. 2019). We selected three simulated datasets (DREAM, sim50, and sim101; Figure 4A-C; Supplementary Table 4) and three cancer datasets (BRCA, melanoma-CML and NTRK; Figure 4D-F; Supplementary Table 5) for this comparison. We analyzed the six datasets using each of the callers with default parameters, then calculated the precision, recall, and F1 scores of each caller. F1 scores measure the overall accuracy by examining the relationship between true positives and false positives, and were calculated using the standard F1 score formula as the harmonic mean of the precision and recall. Caller outputs were then run jointly through MetaFusion and precision, recall, and F1 scores of these results, containing fusions identified by at least two callers each, were calculated as well.

**Figure 4.**
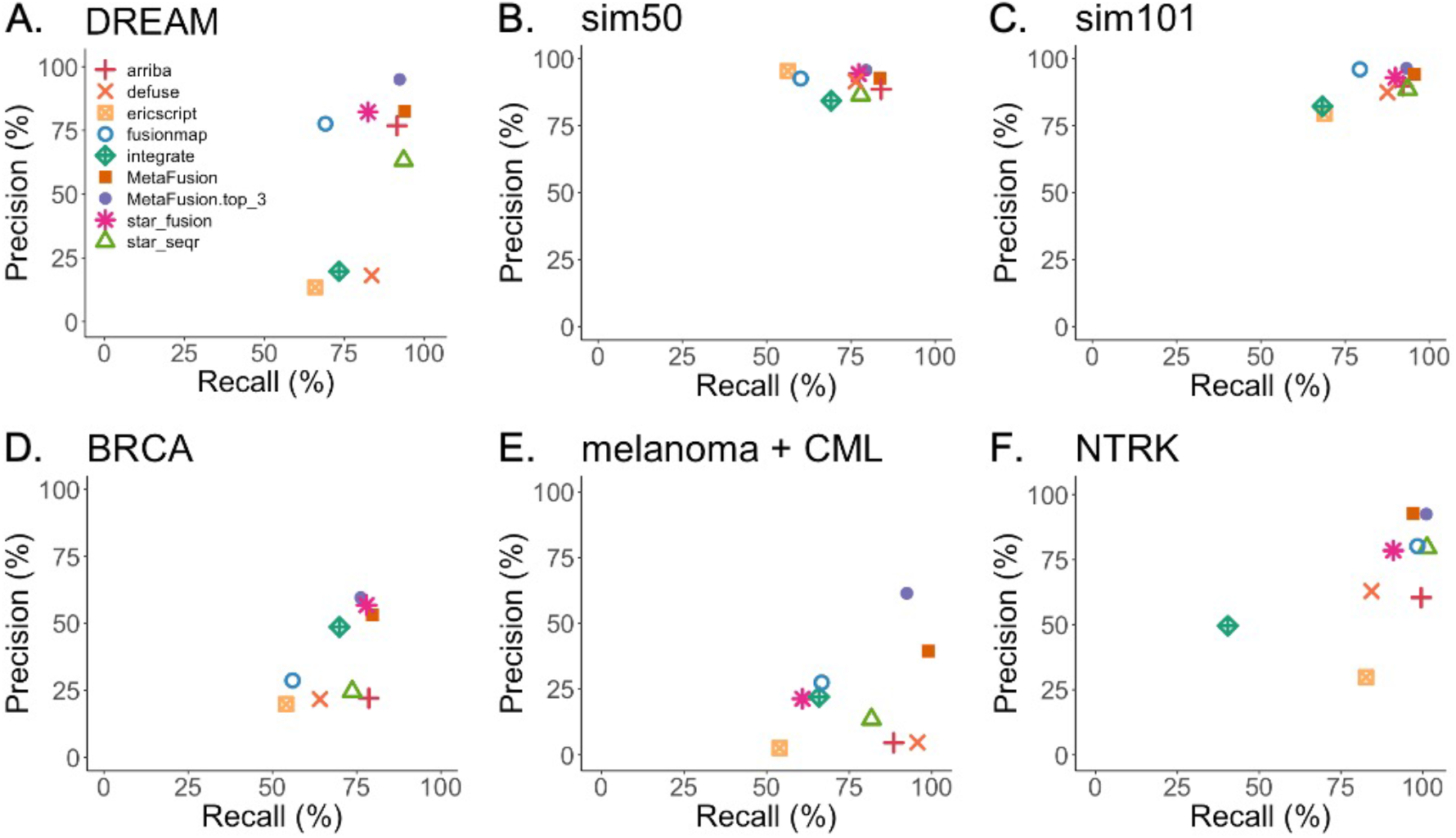
MetaFusion consistently outperforms individual fusion callers on simulated and real datasets. Precision and recall plots showing performance of seven fusion callers, MetaFusion, and MetaFusion.top_3 (using only STAR-Fusion, Arriba and STAR-SEQR) across six datasets. This includes the simulated **(A)** DREAM, **(B)** sim50 and **(C)** sim101 datasets, and the real **(D)** BRCA, **(E)** melanoma-CML, and **(F)** NTRK datasets. MetaFusion generally outperforms individual callers across all three simulated datasets. It performs comparably to STAR-Fusion on the BRCA dataset and outperforms all callers on the melanoma-CML and NTRK datasets. Generally, both MetaFusion and MetaFusion.top_3 showed improved performance compared to individual callers, however a trade-off between precision and recall is often observed between the two.

Counting false positive calls was done on a per-sample basis. For example, if a false positive fusion is present in three samples, it would be counted as three separate false positives. It should, however, be noted that MetaFusion represents fusions present in multiple samples in a single row in the *final*.*cluster* output file (with the names of all affected samples shown in the corresponding column), hence the number of entries in the final output file may be fewer than the sum of true and false positive calls.

MetaFusion generally outperforms individual tools for all six datasets, as indicated by precision, recall, and F1 measurements (Figure 4; Supplementary Table 4, 5). This is also true for MetaFusion.top_3. A trade-off does, however, exist between recall and precision; inclusion of all seven callers results in improved recall at the cost of precision, as MetaFusion.top_3 consistently reports fewer true positives and false positives than MetaFusion, with the exception of the NTRK dataset, where both iterations produce the same number of true positive and false positive calls. Nonetheless, individual callers still typically report fewer true positives than either iteration of MetaFusion. Moreover, in most instances MetaFusion and MetaFusion.top_3 have similar F1 scores, indicating comparable performances. This suggests that ensemble approaches will generally lead to improved results, however users may want to consider their choice of fusion callers, depending on their research needs and whether they would prefer to prioritize precision or recall.

MetaFusion performs favourably even though our combination of seven callers included tools with generally inferior performance on both real and simulated datasets. This is because false positive calls tend to be uncorrelated among methods, and are removed by MetaFusion’s filters and joint calling approach. Thus, MetaFusion can be used to improve upon callers with lower performance to provide high confidence candidate fusions.

In instances where either the precision or the recall of an individual caller is greater than that of MetaFusion, MetaFusion’s F1 score is often higher, indicating better overall performance. For example, both EricScript and STAR-Fusion have greater precision than MetaFusion for the sim50 dataset, yet MetaFusion has a superior F1 score due to its comparatively improved recall (Supplementary Table 4).

Importantly, MetaFusion outperforms individual fusion callers on all three real datasets (Figure 4 D-F, Supplementary Table 5). This is likewise true for MetaFusion.top_3. It should be noted that fusion calling tools generally have poorer performance on real datasets compared to simulated data. This trend remains true with both iterations of MetaFusion, as the pipeline relies on the final results of these tools for its input. Furthermore, complete truth sets for real cancer data cannot be known with certainty, and are often amalgamated from various sources in the literature (Asmann et al. 2011; Edgren et al. 2011; Kangaspeska et al. 2012; Maher et al. 2009). It is thus possible, and even likely, that fusions labelled as false positives in these samples are not yet part of the known truth set (Haas et al. 2019). This may explain why the precision of MetaFusion is lower for the BRCA and melanoma-CML datasets compared to simulated data, and why a higher ratio of false to true positives is detected in the melanoma-CML dataset.

### 3.2 Negative control benchmarking with BEERS dataset

Continuing our benchmarking, we used the BEERS negative control dataset, which contains no true positives, to compare how false positives are reported by MetaFusion and MetaFusion.top_3 compared to individual callers. MetaFusion.top_3 showed the best performance, detecting a single false positive (Figure 5). MetaFusion was only outperformed by FusionMap, which identified four false positives compared to MetaFusion’s five. Callers such as deFuse and EricScript identified over 200 false positives each (275 and 258, respectively). Interestingly, only one false positive fusion was identified by all seven callers, further highlighting the difference in fusion calling approaches across these tools. MetaFusion is able to account for these differences by requiring a minimum of two callers to identify any given fusion, which in turn vastly improves upon filtering out false positives compared to most individual callers.

**Figure 5.**
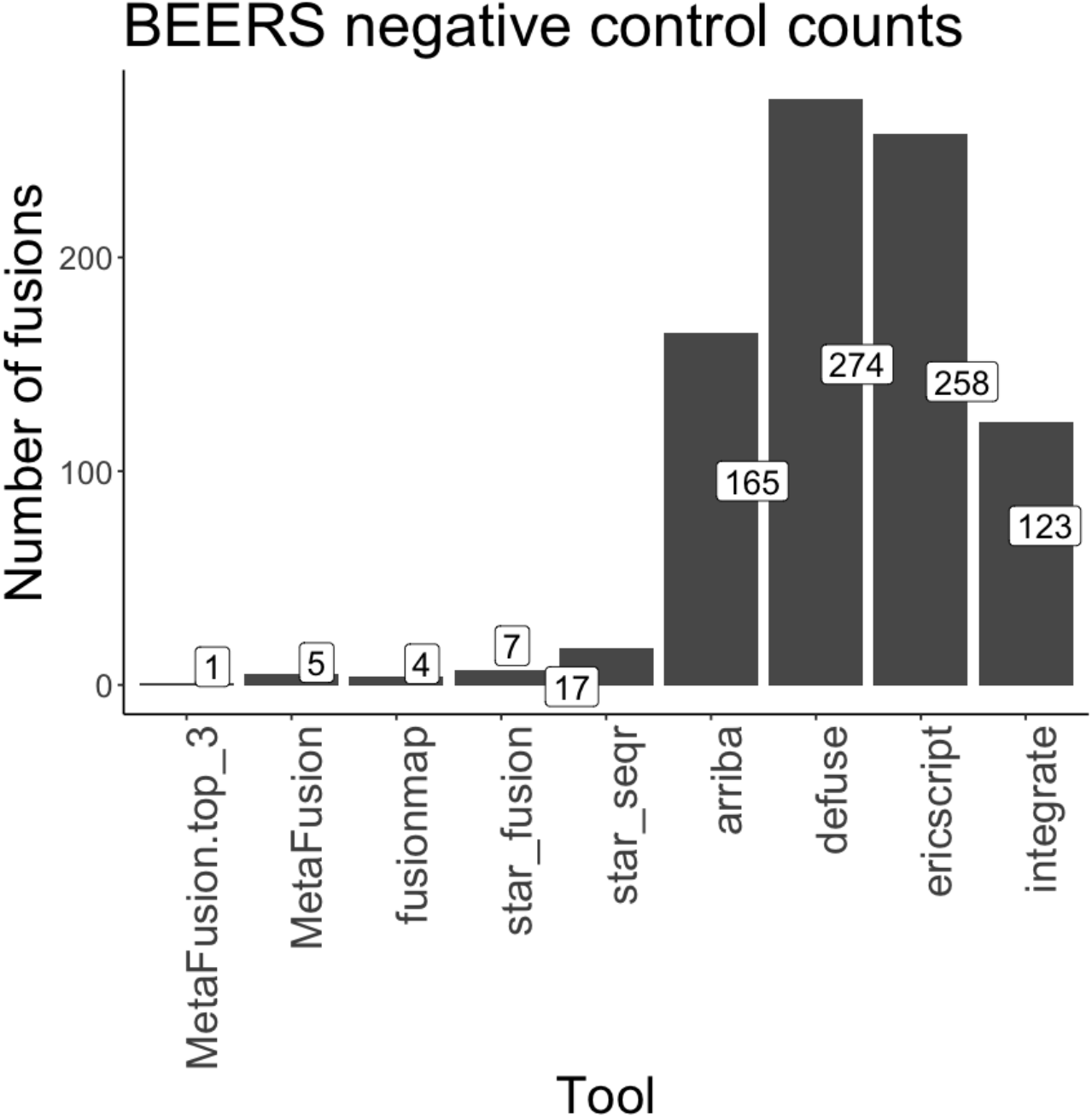
MetaFusion has a low false positive detection rate compared to most individual callers. The BEERS negative control dataset was used to evaluate the detection of false positive calls by MetaFusion, MetaFusion.top_3 and seven individual callers. MetaFusion detected five false positives, outperforming all callers except FusionMap, and MetaFusion.top_3 detected only 1 false positive. Most callers identified a significantly larger number of false positives, particularly deFuse and EricScript.

We also noted that all false positives identified with MetaFusion belonged to the CodingFusion category. FusionMap, STAR-Fusion, and STAR-SEQR, which all performed similarly to MetaFusion, found false positive fusions in a broader range of categories. Nonetheless, CodingFusions tended to predominate, comprising three of the four false positive fusions identified by FusionMap and eight of the 17 identified by STAR-SEQR. STAR-Fusion identified only two CodingFusion false positives. It is likely that this category was so strongly represented because CodingFusions are prioritized by fusion callers due to their relevance in cancer research.

### 3.3 Identifying *cis-SAGe* fusions

RNA fusions can occur due to alternate mechanisms such as trans-splicing or cis-splicing of adjacent genes (cis-SAGe), in which neighbouring genes are transcribed into a single pre-mRNA (Qin et al. 2015). Cis-SAGe fusions such as ReadThroughs are often considered a part of normal biology or transcriptional noise (Babiceanu et al. 2016; Tang et al. 2017) and many tools remove them, prioritizing CodingFusions. Yet some cis-SAGe fusions are translated into fusion proteins, and may occur uniquely in certain types of cancer and serve as disease biomarkers (Varley et al. 2014; Qin et al. 2014, 2016; Rickman et al. 2009). Instead of discarding these fusion calls, MetaFusion stores them in a separate *cis-SAGe*.*cluster* file that can be used for downstream analysis (see Methods, Figure 1). Further details about this are available in the Supplementary Results and Supplementary Figure 1.

### 3.4 MetaFusion uses FusionAnnotator to identify cancer-related fusions in databases

Once the final output of candidate fusions has been obtained, it can be difficult to determine what should be prioritized for further investigation. Cross-referencing oncogenic fusion databases can identify calls that have been previously validated in other forms of cancer, and may thus be of more interest for downstream analysis. FusionAnnotator, which is an optional feature of MetaFusion, leverages such databases and can be used on .*cluster* output files to enrich for cancer-related fusions. This is done based on gene name.

For example, MetaFusion provides 76 total calls for the BRCA dataset, 58 (76%) of which are in cancer fusion databases as indicated by FusionAnnotator. 41/42 (98%) BRCA true positive calls identified with MetaFusion are among these 58. The remaining 17/58 fusions may thus also contain true positives that were not validated when this truth set was established, making them strong candidates for further experimental analysis.

It should be noted that some cancer fusion databases do contain certain fusions found in normal tissues (Singh et al. 2020). Therefore, although enrichment for hits using FusionAnnotator is useful in identifying and prioritizing fusions that may be expressed in cancer samples, these fusions are not always cancer-specific.

### 3.5 Ranking MetaFusion calls by number of callers

Using multiple tools is the best practice in various fields of genomics, such as SNV calling (Fang et al. 2015; Huang et al. 2019) and fusion calling. This was demonstrated by a study in which 23 callers were used to examine the same BRCA dataset used here, with results showing that implementing three or more callers improved fusion detection (Haas et al. 2019). Specifically, increasing fusion caller number led to enrichment of true fusions that have been experimentally validated.

Likewise, assigning a rank to MetaFusion calls based solely on the number of callers by which they are identified highly correlates with true fusion calls in benchmarking datasets (Figure 6; Supplementary Table 6). Indeed, using *callerfilter7* on MetaFusion output results in 100% precision for all benchmarking datasets (Figure 6). This ranking system is particularly meaningful for the real BRCA dataset, where 14/19 calls made by five to six callers and all 21 calls made by seven callers are experimentally validated true positives.

**Figure 6.**
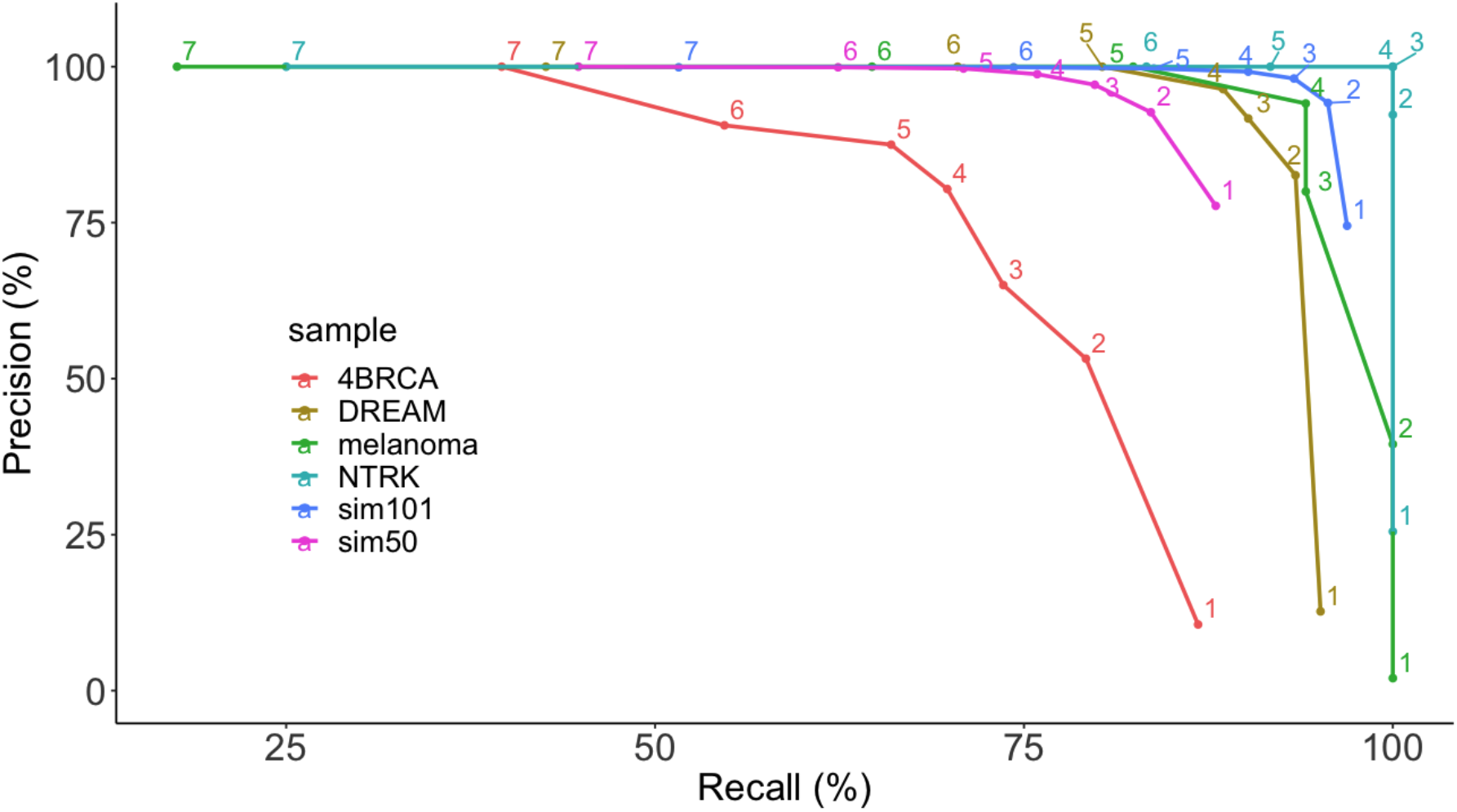
Precision-recall curves for fusions identified by one to seven fusion callers. Precision and recall were evaluated for the six benchmarking datasets (DREAM, sim50, sim101, BRCA, melanoma-CML and NTRK), for fusions identified by an increasingly stringent number of callers. As the caller requirement increased, recall decreased but precision rose. In all six datasets, fusions identified by all 7 callers were true positives.

Although recall decreases as the number of required callers increases, precision can improve substantially. Our results thus suggest that fusions detected by multiple callers are indeed more likely to be true fusions expressed in a given sample. Additionally, consistently higher improvement in precision was seen when going from one caller to two callers. Fusion calls in the MetaFusion *final*.*cluster* output file are therefore sorted by the number of callers that identify them, in descending order.

## 4. Discussion

Gene fusions have well-documented oncogenic effects, making their precise detection critical for both clinical and research applications. Currently available fusion callers, however, may have poor precision or recall, give ambiguous outputs that are difficult to parse, and provide inconsistent results between tools. Here we introduce MetaFusion, a tool for consolidating and prioritizing fusion calls from multiple callers. The MetaFusion pipeline leverages the recall of chosen callers, standardizes format and gene naming between callers, merges the fusion calls, and implements a series of stringent filters to provide a concise final output of fusion candidates. It consistently outperforms individual callers, overcoming limitations of current approaches including high false positive rates, poor recall, lack of a common output format, and inconsistent gene naming. MetaFusion is also equipped with components that allow for further benchmarking and database cross-referencing, making it a valuable tool for cancer and genetics research.

Key to its usability is that MetaFusion is a caller-agnostic tool, meaning that it can be used with any number and combination of upstream fusion calling tools. Our work shows that various iterations of MetaFusion will generally out-perform individual callers, regardless of the upstream callers that have been included (Figure 4, Supplementary Figure 1, Supplementary Table 4, 5). While a trade-off between precision and recall is present (Figures 4, Figure 6), this is not inherently unfavourable. In instances where higher confidence in a smaller number of calls is desired, caller combinations yielding higher precision would be preferred. In contrast, researchers looking to obtain as wide a breadth of candidate fusions as possible may prefer caller combinations yielding higher recall. We also show how selecting callers that are particularly efficient with identifying certain types of fusions, such as cis-SAGe fusions, can impact results (Supplementary Figure 1).

A hallmark of MetaFusion is the seamless integration of multiple fusion callers with the standardization of individual caller outputs into CFF. While various fusion calling tools are readily available, they can vary significantly in the formatting and content of their outputs, making consolidation and direct comparison of their results difficult. Furthermore, each fusion caller has a unique method of identifying fusions such that not all fusions will be identified by all callers, causing the combining of caller outputs cohesively to remain a challenge. We created the CFF as a means to unify the file formats of each tool used for fusion analysis. This allows for any combination of fusion callers to be used with MetaFusion. Users may choose any callers available to them as input into the pipeline, as opposed to being limited to a specific set of tools, such as with other ensemble-based fusion calling methods. Moreover, MetaFusion will only improve as callers with more precise fusion detection capabilities are developed. For these reasons, we hope that future fusion calling tools will include CFF as an output format for their pipelines.

Merging results from different fusion callers is a non-trivial task, and must be done carefully to generate a combined output that is concise and complete. MetaFusion merges based on both gene names and breakpoints, in contrast to other joint calling approaches that rely on one of these two components (Beccuti et al. 2014). To accurately merge fusions based on gene names, MetaFusion renames genes prior to merging, ensuring consistent gene naming across all fusions. To our knowledge, MetaFusion is the only software which provides caller-agnostic, publicly available, robust merge functionality for the integration of fusion predictions.

Our graph clustering method of merging allows for multiple points of contact between groups of similar calls, reducing the chance that a matching call will be orphaned. Calls can be merged together even if they do not intersect directly (Figure 3). This eliminates the need for manual result curation to check for orphaned calls that may have escaped merge, increasing merging reliability. When performing large-scale analyses on hundreds of samples, robust merge is especially important, as large and ambiguous output files are arduous to parse, can delay biological findings, and cause important fusions to be missed.

MetaFusion offers novel functionality that can accelerate biological research by providing researchers and clinicians with an output file which is concise, complete, prioritized, and stringently filtered. MetaFusion’s filters take advantage of the upstream custom annotation and robust merge to remove false positives and prioritize fusion results. Removing fusions called by fewer than *N* callers (*callerfilterN*), filtering out fusion categories enriched for false positives (ReadThrough, AdjacentNoncoding) (Figure 2), and subsequent ordering by number of callers allows for a high-confidence and organized output file. The Benchmarking Toolkit also allows for easy evaluation of new caller combinations using the truth sets provided along with the MetaFusion software. This is a unique feature, as most fusion calling tools do not provide built-in benchmarking functionality. Once caller outputs have been merged, a series of filters is used to refine the results. Enrichment for cis-SAGe RNAs – which most callers discard – or for fusions found in cancer databases allows for further tailoring of the MetaFusion pipeline to specific research questions. Additionally, MetaFusion’s category annotation (Figure 2) allows the user to filter their results by fusion characteristics or identify the types of fusions more likely to be called by specific methods.

While many studies rely on a single fusion calling tool for analysis, we show that a joint fusion calling approach with robust merge and filtering yields improved results. Individual callers may perform inconsistently, have subpar precision and recall, and produce large and ambiguous result files that can impede insight into genetic drivers of disease. MetaFusion offers a novel approach with a caller-agnostic framework to provide concise and complete results to researchers and clinicians for identification of gene fusions in cancer. Future directions for MetaFusion include features for use in the clinic, including addition of annotations for retained and removed protein domains, integration of caller frame information, separation of calls per-sample and per-breakpoint, and addition of an SQLite database for clinical use to track historical calls. These features will add to the flexibility and utility of MetaFusion, making identification of oncogenic fusions in both research and clinical contexts easier and more precise.

## 5. Software availability and implementation

MetaFusion is a free software tool implemented in Python and R, with bash scripts used as wrappers. The MetaFusion source code is available on GitHub at [https://github.com/ccmbioinfo/MetaFusion]. For convenience and ease of installation, a platform-independent Docker image containing installed dependencies is available at [https://hub.docker.com/r/mapostolides/metafusion]. Reference files to run MetaFusion can be downloaded from figshare at [https://figshare.com/articles/dataset/Metafusion_reference_files/12855080] and [https://figshare.com/articles/dataset/FusionAnnotator_required_files/12915455]Instructions for downloading the Docker container, running MetaFusion software, discerning output files and fastq file data access can be found at https://github.com/ccmbioinfo/MetaFusion/wiki. The individual fusion callers used by MetaFusion are available at their respective software repositories (see References).

## 6. Supplementary Material

Supplementary material provided online.

## 7. Acknowledgements

This work was supported by the Canadian Center for Computational Genomics (C3G), part of the Genome Technology Platform (GTP), funded by Genome Canada through Genome Quebec and Ontario Genomics. The authors would like to thank Robert Eveleigh for his help with benchmarking, acquisition of the DREAM dataset, and Genpipes implementation of fusion callers, Brian J Haas for his assistance and valuable feedback on this study, and Man Yu for help with data preparation.

