## Supplementary Materials for "MetaFusion: A high-confidence metacaller for filtering and prioritizing RNA-seq gene fusion candidates"

^1^Centre for Computational Medicine, The Hospital For Sick Children, Toronto, ON, Canada; ^2^The Arthur and Sonia Labatt Brain Tumour Research Centre, The Hospital for Sick Children, Toronto, ON, Canada; ^3^Department of Laboratory Medicine and Pathobiology, University of Toronto, Toronto, ON, Canada. ^4^Division of Pathology, The Hospital for Sick Children, Toronto, ON, Canada. ^5^Genetics and Genome Biology Program, The Hospital for Sick Children, Toronto, ON, Canada. ^6^Department of Computer Science, University of Toronto, Toronto, ON, Canada. ^7^University Health Network, Toronto, ON, Canada.

### **Supplementary Methods**

**Merge details**

Merging is done in four steps, outlined below.

First, intersection edges are generated using bedtools pairToPair function (Quinlan and Hall, 2010). A BED file is generated in which every line is a separate fusion call. The file is then intersected with itself, and an edge is generated from each intersection. Breakpoints must be within 100 base-pairs of one another to be considered a match, since this was the distance we found consolidated most relatively close breakpoints. An example of edges generated based on breakpoints is represented in Figure 3B.

Second, edges are generated by matching calls based on gene names. Names must be matching exactly. A hash table is created which uses head- and tail-gene names as keys, and collects FIDs which match the “Head--Tail” key (i.e. {key: [fusion_id list]}). Then, for each key, an edge is created for each FID pair in its corresponding FID list. An example of edges generated based on gene names is shown in Figure 3C.

Third, a graph is generated using the R package RBGL (Carey *et al.*, 2020). Edges from both breakpoint intersections and gene name matches are combined, and a graph is constructed from the edge set. Since edges contain only FIDs, the graph is built unaware of which edges are due to gene name matches or breakpoint intersections (Figure 3D). For each connected subgraph, a list of FIDs is generated which correspond to CFF file entries.

Fourth, the CFF entries are converted to a cluster format, MetaFusion’s final output format (Supplementary Table 1B). Delimiter-separated lists are generated for fields such as sample, tool, and breakpoint, which may differ among merged calls.

Merging is done based on both gene name and breakpoint information as using either one on its own will create a result file with unmerged similar calls, and will remove calls which would otherwise pass the ensemble tool cutoff.

With respect to breakpoints, the existence of variants and splicing isoforms can result in breakpoints thousands of base pairs away. Thus, relying on breakpoints alone, even with a pairToPair-like tool with a breakpoint range parameter (i.e. -slop), may result in calls from callers with poor breakpoint accuracy not being properly merged, creating a result file containing incorrect fusion predictions. Moreover, fusion splicing isoforms and variants will not be combined with one another, resulting in duplicate calls.

With respect to merging based on gene names alone, a common issue in bioinformatics is gene naming inconsistency between tools and databases. Fusion callers often use their own custom databases, and may not get updated regularly (either by the user, the developer of the tool, or both). For this reason, renaming is necessary, but not sufficient, for merging fusion call results. Using only gene names prevents merging of genes that were unable to be renamed correctly. Furthermore, callers may predict different RNA species with identical fusion breakpoints (Figure 3C), which will not be merged despite being the same event. Both of these situations result in a file containing unmerged calls, and may remove calls which do not pass the “--num_tools” parameter threshold.

**Filter descriptions**

*Blocklist filter*. MetaFusion’s *blocklist* filter is based on part of Arriba’s blacklist file. This filter is used to block out known ReadThroughs, T-cell receptors, MHC complexes and immunoglobulins, as fusions involving these genes are less frequently of interest in cancer-specific research. This filter intersects the blacklist file’s “recurrent breakpoints”, “T-cell receptors”, and “MHC complexes” paired regions with fusion breakpoints using bedtools pairToPair. Those which intersect are removed from the final output. Users can include a personal list of false positive fusions to this file if they wish to remove them from their results.

*CallerfilterN*. This filter is used to remove fusions identified by fewer than *N* number of fusion callers. Fusions called by *N* or more callers are kept as part of the final output. The user is able to set the value of *N* to their desired threshold. Unless otherwise indicated, for this study we use *callerfilter2* (i.e. calls have to be made by at least two callers).

*ReadThrough filter*. ReadThrough fusions, or those consisting of two adjacent coding genes in which the head gene is immediately upstream of the tail gene, occur in healthy tissues (Babiceanu *et al.*, 2016) and are more likely to be the result of cis-splicing than chromosomal rearrangements (Qin *et al.*, 2015; Tang *et al.*, 2017), making them seldom of interest in cancer research. As such, MetaFusion includes a *ReadThrough* filter, which removes entries categorized as ReadThrough fusions during reannotation. If they are of interest, ReadThrough fusions can still be viewed in a separate *cis-SAGe.cluster* file, decribed below.

*Adjacent noncoding filter*. The *adjacent noncoding* filter removes all fusions in the TruncatedCoding, TruncatedNonCoding, and NoHeadGene categories whose constituent gene partners are within 100kb of one another, as we noticed that fusions with these characteristics are heavily represented among our negative control dataset (Supplementary Figure 2). Using this filter also removed false positives from our benchmarking datasets and did not affect true positive counts. These fusions are included in the *cis-SAGe.cluster* file described below. Since our negative control is designed to contain no fusions caused by chromosomal rearrangements (further detail in Datasets and Results sections), we developed this filter.

**cis-SAGe.cluster file**

MetaFusion stores cis-SAGe fusions, such as ReadThroughs, in a separate *cis-SAGe.cluster* file, instead of simply discarding them. Not all cis-SAGe fusions are necessarily ReadThroughs, therefore this file also contains SameGene fusions as well as those flagged by the *adjacent noncoding* filter, as their breakpoint proximity and orientation may be indicative of cis-splicing. Fusions in the *cis-SAGe.cluster* file are removed from the final.cluster output using the above-described filters.

#### **Supplementary Results**

**cis-SAGe fusions**

Fusions such *SCNN1A-TNFRSF1A* and *CTSD-IFITM10* are translated into fusion proteins, and can contribute to cancer progression (Varley *et al.*, 2014). Both *SCNN1A-TNFRSF1A* and *CTSD-IFITM10* are identified by MetaFusion in our BRCA data. Additionally, *SLC45A3-ELK4*, a ReadThrough present in the urine of men at risk for prostate cancer (Rickman *et al.*, 2009), is detected in our prostate data. These cis-SAGe fusions are filtered out of the final.cluster results by MetaFusion’s filters, as they cannot be distinguished from cis-SAGe RNAs found in normal biology, and are instead stored in the datasets’ respective *cis-SAGe.cluster* files.

We further show that *cis-SAGe.cluster* files are enriched for cis-SAGe fusions (Supplementary Figure 1) by analyzing a prostate cancer dataset containing 44 cis-SAGe fusions verified by Sanger sequencing (Qin *et al.*, 2015; Kumar *et al.*, 2016). It should be noted that the prostate dataset is not used as a benchmarking dataset, nor should it be. Benchmarking of the MetaFusion tool is based on curated truth sets of known fusions occurring between non-adjacent genes. Since ReadThrough fusions are usually transcriptional noise, their aberrant expression is a separate research question from studying fusions caused by chromosomal rearrangements.

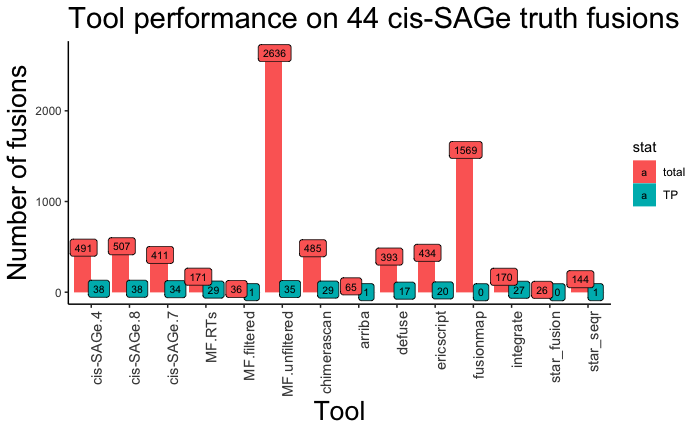

**Supplementary Figure 1: Investigation of cis-SAGe fusions with MetaFusion and various individual caller configurations.** For investigation of cis-SAGe RNAs, the *cis-SAGe.cluster* file can be used. Improved recall can be seen with the addition of chimerascan, a caller which detects cis-SAGe RNAs. Only 4 callers are needed to obtain optimal recall. (cis-SAGe.4: the cis-SAGe output file for MetaFusion outputs using the 4 callers that identified the most cis-SAGe true positive fusions - ChimeraScan, INTEGRATE, EricScript, and deFuse; cis-SAGe.8: cis-SAGe outpit file with chimerascan in addition to the 7 standard callers we used upstream of MetaFusion; cis-SAGe.7: cis-SAGe output file using the seven standard callers we used upstream of MetaFusion, MF.RTs: ReadThrough fusions identified by MetaFusion; MF.filtered: usual output of MetaFusion, which is not useful when looking at cis-SAGe RNAs; MF.unfiltered: merged metafusion output from all 7 callers, before filters are applied; TP: true positive)

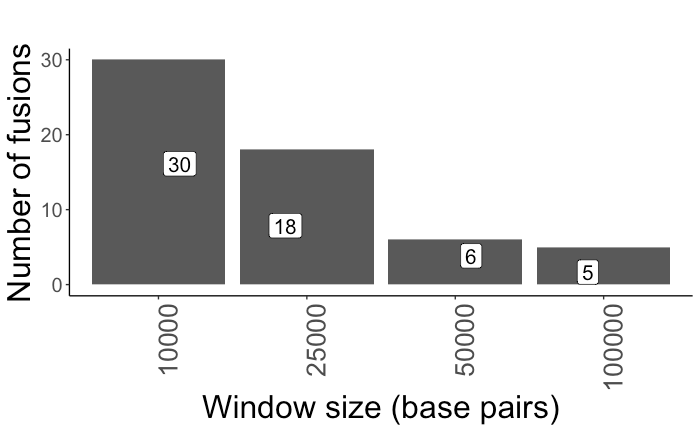

**Supplementary Figure 2: Investigation of various window sizes for the Adjacent noncoding filter.** The window size of 100kb was found to be ideal for removal of all Adjacent noncoding fusions.
